# Peri-somatic modulation of diffracted light and its variation with consciousness

**DOI:** 10.1101/2025.02.03.635129

**Authors:** Santosh A. Helekar

## Abstract

Existing noninvasive neurophysiological recording methods, such as electroencephalography, magnetoencephalography and functional magnetic resonance imaging, can acquire neural correlates of consciousness, but they cannot mechanistically differentiate between conscious and unconscious states. Here we describe a method that detects a new physical effect of apparent neural origin in the peri-somatic space using a newly developed instrument and test whether it might allow such differentiation. This effect is a slow decrease in the intensity of diffracted low power laser light emitted within a photoelectronic instrument placed close to the body of an awake subject. It is markedly attenuated by general anesthesia in mice and unconsciousness in patients. Its direction is reversed when the instrument is exposed to live invertebrates or to the decapitated head of a euthanized mouse. We provide evidence to rule out all plausible nonspecific physical and non-biological explanations for the photo-modulatory effect in our estimation, such as those arising from thermodynamic and electromagnetic influences. We have also been able to record it in a near-complete vacuum. These findings point to the existence of a previously unrecognized biophysical phenomenon that appears to be related to a mechanism underlying consciousness.

## Introduction

The existing methods of studying brain function rely on our understanding that neural information processing involves action potentials and synaptic transmission occurring within neuronal networks^1^. These methods have greatly expanded neuroscientific knowledge of the underlying mechanisms of sensory processing, object representation, self and body image representation, processing of emotions, interoception and proprioception, memory storage and retrieval, volition and motor planning and action ^2–11^. However, they cannot yet clearly distinguish between conscious and unconscious brain activity or detect a neural process that may be uniquely related to consciousness in any organism^12–14^. The phenomenon of consciousness itself continues to elude a scientific explanation, despite having been the subject of intense philosophical and scientific debate for centuries^12–14^. Recently, a large-scale experimental effort has been undertaken to test two prominent theories of consciousness, Integrated Information Theory and Global Neuronal Workspace Theory, using functional magnetic resonance imaging, magnetoencephalography, and intraoperative electrophysiological recordings under an adversarial collaboration^15^.

Neuroscientific investigations focus on the notions that neuronal electrical activity, computation in neural networks, mode of distribution of neural information, complexity of information processing, and/or predictive processing, are somehow responsible for conscious awareness or subjective experience^16^. Nonetheless, the limited progress in pursuing these avenues of research to obtain compelling evidence in support of a singular, most likely explanation for consciousness has led to unconventional speculations involving the possibility that the explanation resides at a subcellular level and involves fundamental physics. For example, there has been a revival of old quantum physical ideas and new proposals inspired by quantum neurobiology because of growing evidence that quantum physicochemical processes play wide-ranging roles in biology, such as quantum superposition and coherence in photosynthesis^20^, inelastic electron tunneling in olfactory ligand-receptor interactions^21^, and weak magnetic field effects on the radical pair mechanism in cryptochromes in avian migration^22^.^17–19^Other quantum effects include light harvesting and exciton diffusion in microtubules which is attenuated by general anesthetics^23^, isotopic effects of xenon as an anesthetic^24^ and increased fluorescence quantum yield in biological networks containing tryptophans^25^..

We have found recently that the intensity of light wave phenomena such as interference and diffraction produced by light from a low power laser light emitting diode (LED) is modulated by proximity to the head of a human subject in a readily detectable and reproducible manner. In this paper we describe for the first time a new noninvasive photoelectronic instrument developed in our laboratory to characterize the above photo-modulatory biophysical effect further, rule out possible known confounding causes for our observations and test whether the modulation is affected by any changes in consciousness due to general anesthesia and cessation of cerebral activity in mice and brain impairment in patients. We report here that, in recordings conducted with this instrument (described below), placement of its sensor in the peri-somatic space for brief durations is associated with a marked distance-dependent change in the intensity of diffracted light, characterized by a decrease in human subjects and mice, and an increase in example invertebrates that were tested. We also observe that the decrease is attenuated by general anesthesia in mice and unconsciousness due to brain injury in patients admitted to the intensive care unit (ICU). Furthermore, a decapitated mouse head shows a paradoxical increase in intensity. These observations suggest a possible relationship of the measured light wave intensity modulation to a putative neural process and offer a potential new method to record and measure a correlate of the relative change in the conscious and unconscious activity of the brain.

## Results

### A noninvasive photoelectronic instrument detects a new peri-cranial physical effect

We developed two versions of a new photoelectronic instrument that measure either interference or diffraction of light waves emitted by a low power laser LED (See Figure 1A showing a schematic diagram of the sensor module and its hypothetical principle of operation, and Figure 1B showing current working prototype used in the experiments described here). For each recording we powered the instrument by connecting it to a laptop computer (Figure 1B) or an Android tablet and recorded the voltage output of four off-center photodiodes that sampled diffracted light in separate channels, using a serial monitor application that logged the data at the sampling rate of 100 Hz. All experiments began by recording the instrumental baseline with no source of activity close to it within a radius of 180 cm for at least 30 min. It took 20 – 30 min for the baseline to stabilize at a constant level. A baseline recording of intensities from 4 off-center channels in an empty room after achieving a stable shows no large amplitude deviations baseline in Figure 1C. A time series of scores of the first principal component of these data are shown in Figure 1D. In the first experiment, after 40 min we instructed each human subject to sit on a chair for 10 – 15 min such that his/her left temple was 1 cm from the sensor module. Light intensity recordings from all channels during exposure of the head to the sensor module consistently showed a large decrease in intensity that peaked at the end of the exposure and declined gradually over ∼30 min (Figure 2A). Principal component analysis of the 4 off-center channel traces to measure the shared variance across all channels showed that >97% of their variance was accounted for by the first principal component. Therefore, we plotted a normalized inverted version of this component as the final representation of the recorded effect in all subsequent experiments. We have expressed the strength of the recorded photo-modulatory response (PR) in arbitrary units (au). Experiments involving transient exposure to the instrument sensor were done in a total of 22 adult humans subjects. 10-min recordings from 7 healthy adult human subjects with the sensor module placed 1 cm from the left temple revealed the strength of PR to be 601.4 ± 61.6 au (mean and standard error of the mean). To rule out any element of subjective bias in the recording of PR and in attributing it to actual exposure of the head to the instrument sensor, we conducted a double-blind experiment in which the subject who was alone in a closed room, sat at random on one of two chairs placed 90 cm apart (a distance at which little or no PR was detected). Each chair was fitted with either an active or an inactive instrument, indistinguishable to the subject. Using an electronic tablet the subject indicated the time and the chair on which he/she sat. The researcher conducting the experiment who was outside the closed room could find out whether the occurrence of the PR matched with the position of the subject only at the end of the experiment. In all recordings conducted in 5 subjects there was a perfect match between the PR and the position of the subject relative to the active instrument (Figure 2B). We then tested the dependence of the PR amplitude on the distance between the sensor module and the head (left temple) by recording PRs at 9 different distances in two experiments, each conducted in 3 subjects. Example recordings from one of these experiments and bar plots of mean amplitudes and standard errors of the mean (SEM) as a function of distance are shown in Figure 2C. The amplitude declines steeply with distance with a half maximal decrease of ∼2.7 cm.

**Figure 1.**
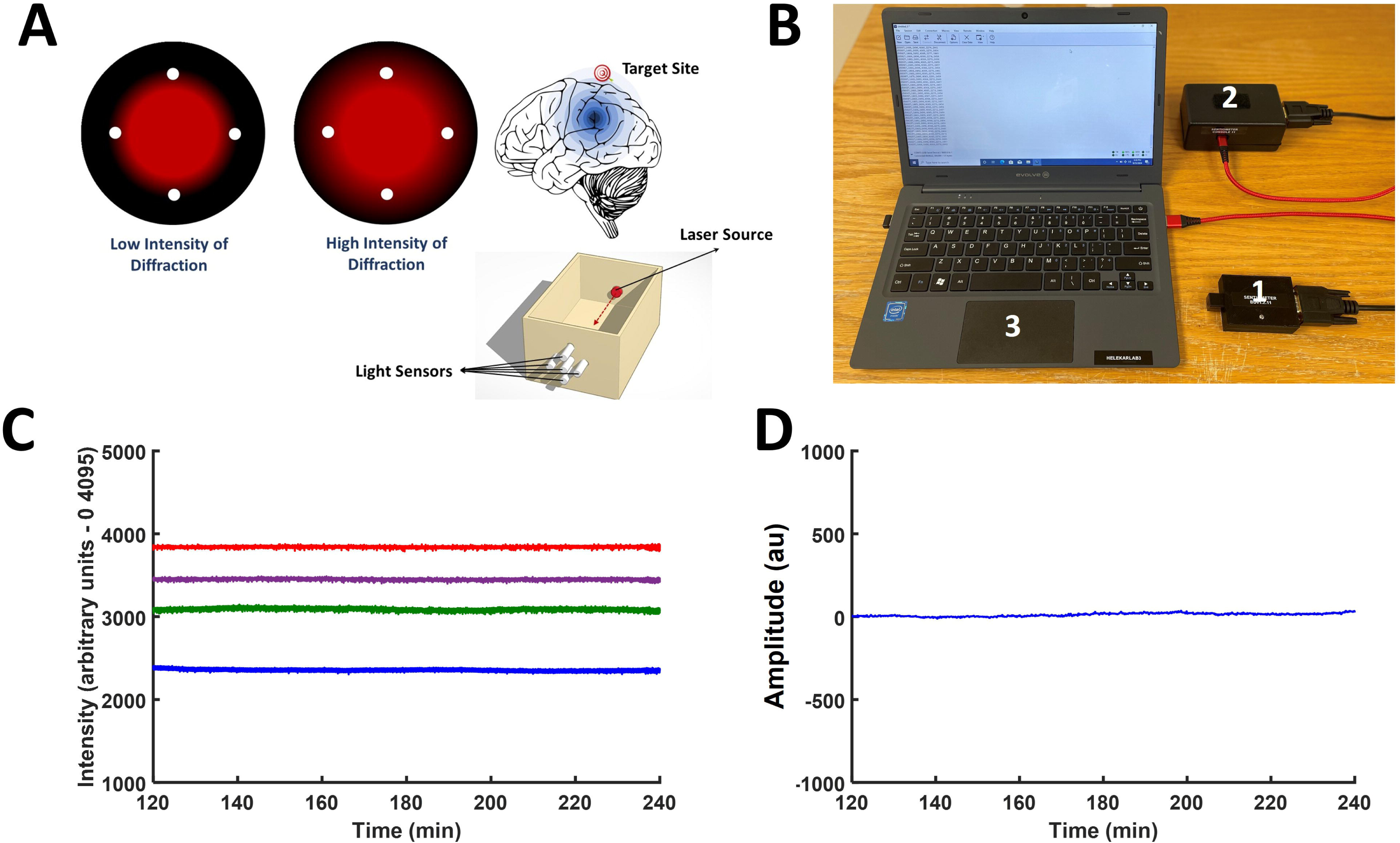
Photoelectronic Instrument Prototype Description and Operation. **A.** A schematic diagram showing the operation of version-2 the photoelectronic instrument. **B.** An image showing the components of the instrument: 1 – sensor module/unit, 2 – controller and 3 – Laptop computer. **C.** Raw sensor output channel recordings in an empty room. **D.** Normalized first principal component score time series derived from **C**.

**Figure 2.**
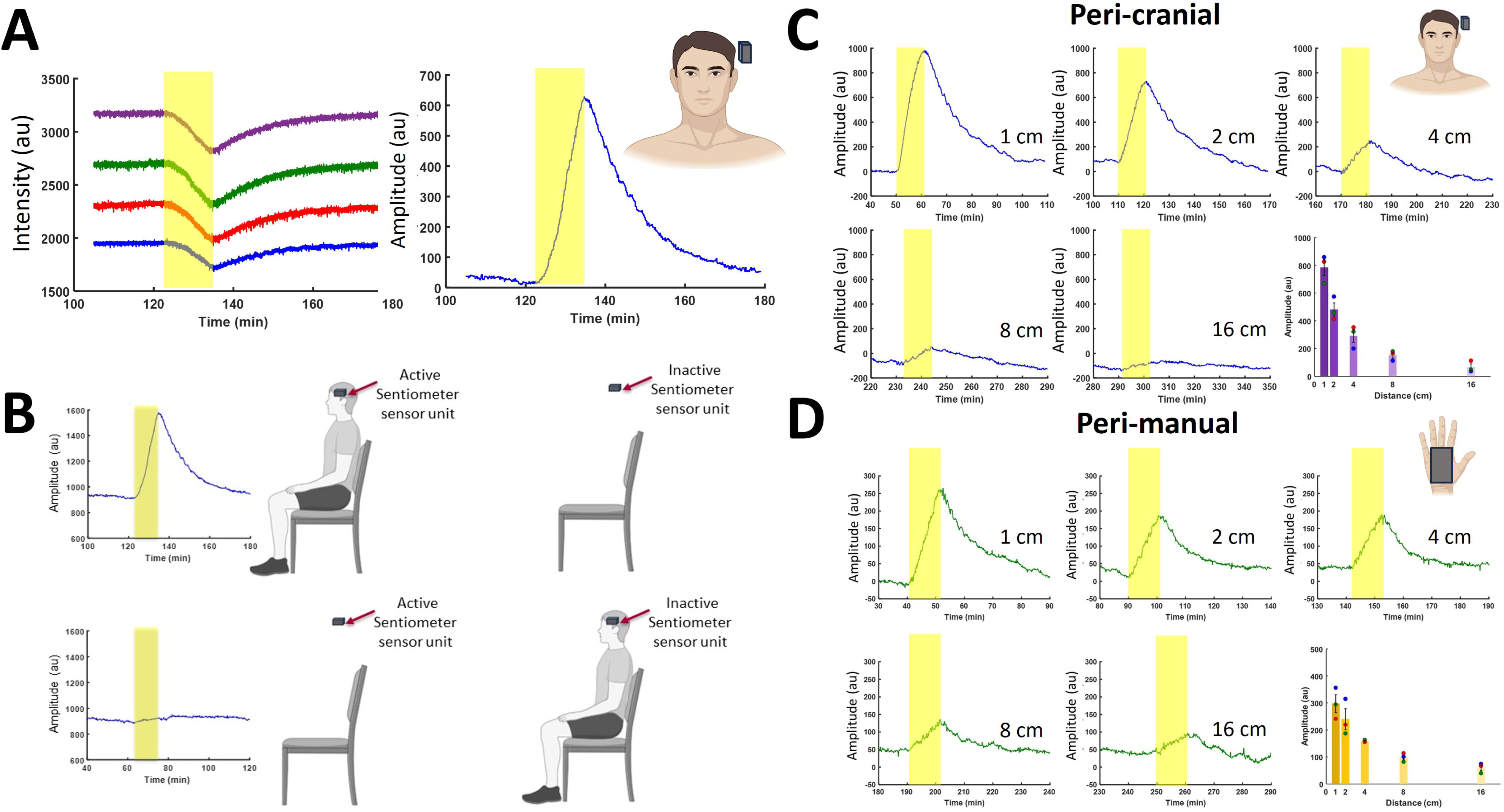
Peri-somatic Photo-modulatory Response (PR) **A.** The light intensity sensor unit when brought close to the head (yellow boxes represent the timing and duration of exposure) in an awake subject produces a robust deviation of the time series away from the baseline with a slow time course of ∼40 min [Left – raw channel recordings, **Right** – normalized first principal component score time series representing the photo-modulatory response (PR)]. Note that the normalization step renders a decrease in diffracted light intensity to be represented as an increase in amplitude of the PR. This convention is used in all figures, below. **B.** A double-blind experiment with active and inactive instruments shows the expected result. **C.** and **D.** PRs are shown varying as a function of distance. The bar plot data is derived from n = 3 each for peri-cranial and peri-manual PRs. The error bars depict standard errors of the mean (SEM) in this and all figures below. Images in the figure are obtained from the stock image database of licensed BioRender software.

### Spread of the physical effect is detected in the peri-somatic space

To investigate whether the measured physical effect can also be recorded close to the rest of the body, we exposed a hand to the sensor module of the instrument at 1, 2, 4, 8 and 16 cm distances. We observed a peri-manual PR (n = 3) that was ∼31%, and significantly (p = 0.014, two-tailed Student’s paired t test, n = 3), reduced in amplitude compared to peri-cranial PR when the two were measured under the same conditions. It also declined with distance like the peri-cranial PR. Traces and bar plots of peri-manual PR are shown in Figure 2D.

### PR is not produced by body heat, humidity, respired air or inaudible sound

Given that temperature affects laser sources and current flow in electronic circuits we tested whether PR is produced by exposure to ∼37°C body temperature. We measured the extent of the increase in temperature produced at 1 and 5 cm from the human head (temporal region). The temperature increases at these distances were 2.3°C and 1.2°C, respectively. The change in the instrumental baseline amplitude due to exposure of a beaker containing water that was warmed to or cooled from ∼37°C in 2°C steps corresponded to 16 au/°C and 13 au/°C for increasing and decreasing temperatures, respectively (see Figure S1). We also conducted recordings at ∼37°C by keeping the sensor at this temperature with a beaker containing warm water (∼37°C water bath) and exposing it to a hand for 30 min at 1 cm. PR shown in Figure 3A is nearly like that at room temperature in amplitude. However, the warmer water bath temperature increased the instrumental baseline. The latter effect requires further investigation. To investigate whether PR is an artifact of movement of air or change in the composition of air due to perspiration and respiration we placed the entire instrument including the connected electronic tablet in a stainless-steel vacuum container (Figure 3B). The vacuum pressure achieved in the container was ∼26 inches of mercury. The peak PR amplitude in response to exposure of the left temple obtained under these conditions was not significantly different from that obtained when the container was left open to air. This result indicates that PR is not due to breathed air or displacement of air produced by inaudible sound or ultrasound waves.

**Figure 3.**
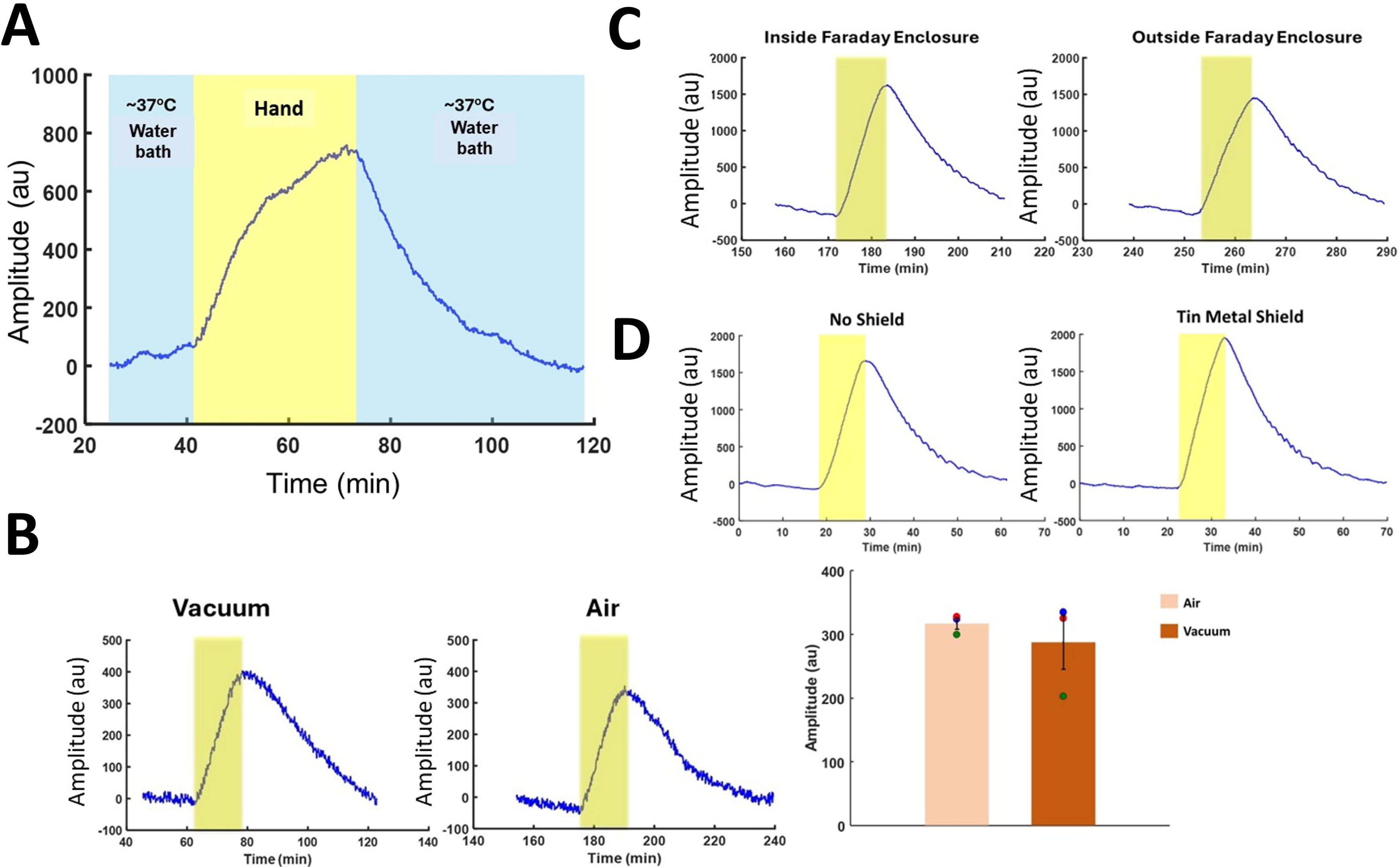
PR is Not Due to Known Physical and Environmental Variables. **A.** Peri-cranial PR recorded with an instrument warmed to ∼37°C (baseline) with a water bath. **B.** Peri-cranial PR recordings at ∼2 cm in a vacuum chamber and in air (n = 3). The colored box represents the timing and duration of exposure as above and in all figures below. The bar plot represents means and SEMs of the PRs under the two conditions. **C.** Peri-cranial PR recordings at ∼1 cm inside and outside a Faraday bag. **D.** Peri-manual PR recordings at ∼1 cm with and without a metal barrier made up of tin (∼0.6 mm thick). Similar results were obtained with aluminum and copper barriers (see *SI Appendix*, Fig. *S2*).

### PR is not due to static and radiating electromagnetic fields

To test whether any perturbation of the static magnetic field was responsible for the observed PR we exposed the instrument sensor to axially magnetized (1.48 T) N52 (0.635 cm diameter and 1.905 cm length) and diametrically magnetized N42 (1.905 cm diameter and 1.905 cm length) cylindrical neodymium magnets at 1 cm distance. At 1 cm the measured magnetic flux was ∼135 mT and ∼152 mT for the two types of magnets, respectively. Recordings showed no deviation of the instrumental baseline in response to the magnets (see Figure S2A). These results rule out the direct influence of a static magnetic field on light intensity recording. The amplitude and time course of the PR produced by the left temple did not differ significantly if the sensor module was placed within a Faraday shield bag or left exposed to radiating radiofrequency electromagnetic fields (Figure 3C). There were also no significant differences in the amplitude and time course of the PR when the sensor module was shielded from the left temple by metal sheets (Figure 3D. tin, Figure S2B copper, and Figure S2C aluminum, n = 3 for each exposure) or left unshielded. These findings rule out any contribution of radiofrequency electromagnetic field interference or static electric fields to the production of PR. Exposing the sensor module to the left temple with the laser beam turned off also did not produce any deviation of the recorded instrumental baseline, indicating that there is no voltage response produced in the photodiodes by any ultraweak photon emissions(33) from the head (see Figure S2D).

### PR is inverted in invertebrates compared to mammals

We asked whether other species of animals produce PR by exposing mice and 4 different invertebrates (9 blue crabs, >200 California blackworms, 3 crayfish and 6 cherrystone clams) to the sensor module. A mouse when exposed to the sensor module for 60 min at <5 cm evoked an PR like that in humans but of a smaller amplitude (Figure 4A). A 60-min exposure in 10 mice produced a peak PR amplitude of 516 ± 32 au. The invertebrates, blue crabs (n = 9) and California blackworms (>200 in a clump) exposed for 60 min to the light intensity sensor produced an inverted PR (Figure 4A). Other invertebrates also produced an inverted response with a similar time course (see Figure S3).

**Figure 4.**
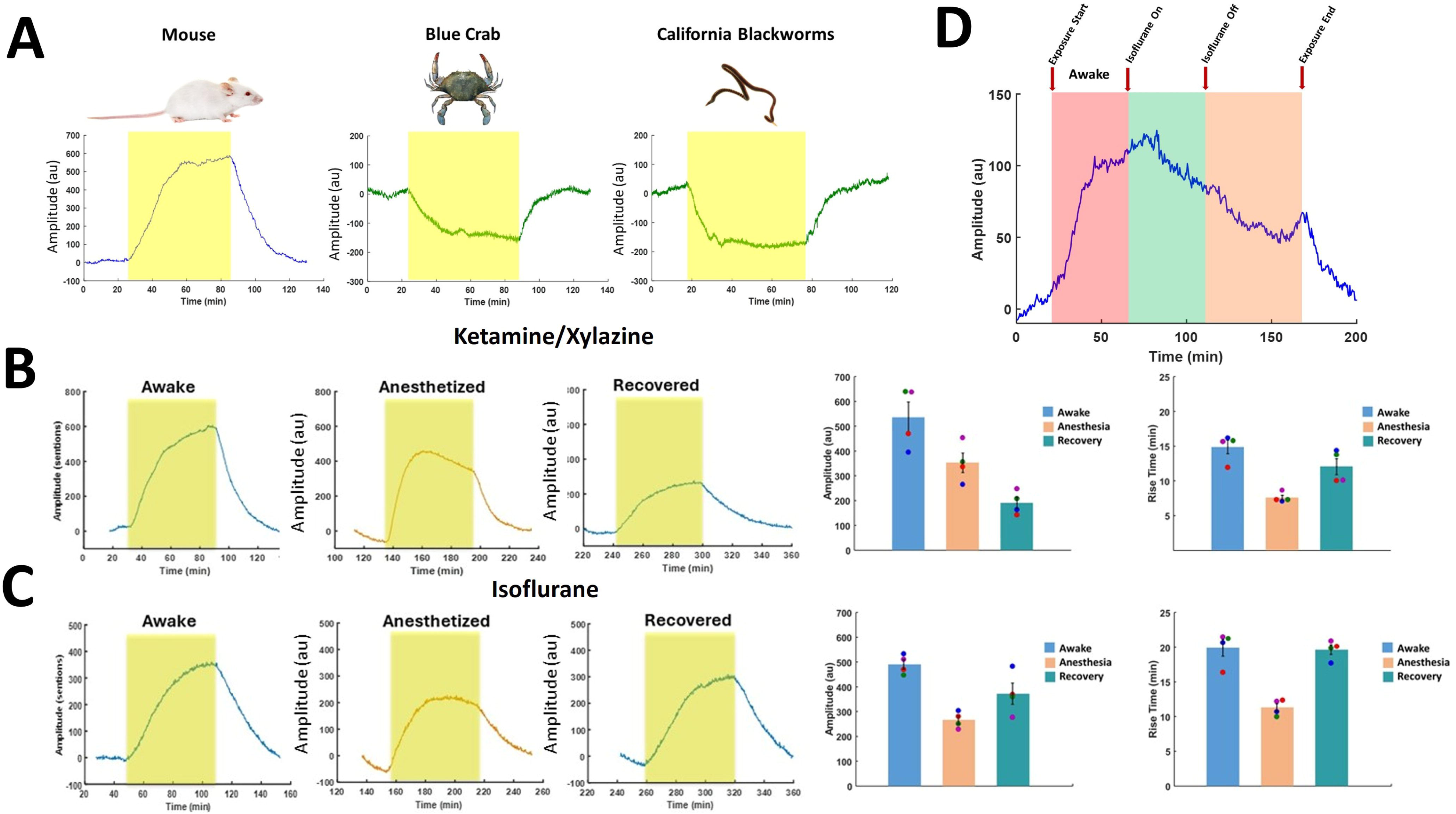
Differences in PR Produced by Other Living Organisms and General Anesthesia. **A.** Peri-somatic PR recordings at ∼1 cm with another mammal, mouse (n = 20) and invertebrates, blue crab (n = 9) and California blackworm (a clump of >200). Mice show upright PRs comparable to humans. Invertebrates show inverted PRs. **B.** Peri-somatic PRs (at <3 cm) before, during and after general anesthesia in mice loosely restrained in a plastic tube for a 60-min exposure. Ketamine and xylazine combination was administered intraperitoneally at 100 mg/kg and 18 mg/kg, respectively (n = 4). **C.** Isoflurane gas was administered at 2% in a small animal anesthesia chamber (n = 3). The bar plots depict means and SEMs of 60-min amplitude and half maximal rise times corresponding to the 3 conditions. Recovery from anesthesia is better seen in half maximal rise time, and not in amplitude. **D.** Continuous recording of the PR from a mouse exposed to the sensor at ∼5 cm before (∼55 min, pink box), during (∼55 min, green box) and after Isoflurane (∼70 min, orange box) general anesthesia. The arrows mark the start and end times of exposure and the on and off times of isoflurane flow into the anesthesia chamber. Images in the figure are obtained from the stock image database of licensed BioRender software.

### General anesthesia decreases the amplitude and shortens the rise time of PR in mice

Substantial evidence indicates that general anesthesia induces complete loss of consciousness^27^. If PR is produced by a mechanism that underlies, or is related to, consciousness, then all general anesthetics should significantly attenuate it. We tested this prediction with an inhalant anesthetic, isoflurane (If) and 3 injectable formulations – ketamine/xylazine (KX), pentobarbital (Pb), and propofol (Pf), administered intraperitoneally in a total of 30 mice. All anesthetics produced similar significant alterations in the time course and peak amplitude of the PR. The mean half-maximal rise time was ∼44% shorter under anesthesia (7.6 ± 0.4 min, n = 4 for KX; 11.3 ± 0.6 min, n = 4 for If) than in the pre-anesthesia waking state (14.9 ± 1.0 min for KX; Figure 4B; 19.9 ± 1.2 min, n = 4 for If; Figure 4C). The mean amplitudes at 60 min were ∼42% lower under anesthesia (353 ± 61 au, n = 4 for KX; Figure 4B; 266 ± 16 au, n = 4 for If; Figure 4C) than before anesthesia (535 ± 39 au, n = 4 for KX; Figure 4B; 490 ± 19, n = 4 for If; Figure 4B). These differences were statistically significant for both the rise time (p = 0.005, n = 4 for KX; p = 0.012, Student’s paired t test, n = 4 for If) and amplitude (p = 0.014, n = 4 for KX; p = 0.0018, Student’s paired t test, n = 4 for If). These effects reversed with recovery except for the decrease in amplitude with KX over the duration of the experiment, likely due to its prolonged blocking action on NMDA receptors^28^. The decreases in amplitude at 60 min and rise time showed small differences in values between the four types of anesthetics (see Figure S3B for Pb and Pf effects). We also carried out continuous light intensity recordings in mice (n = 3) before (∼55 min), during (∼55 min) and after (∼70 min) administration of If general anesthesia to monitor the transition from the awake state to the anesthetized state and the recovery from the latter state to normal wakefulness over a period of 3 hours. Each mouse was kept in a loose restraining tube inside the anesthesia chamber supplied with O_2_ and CO_2_ or air throughout the recording. The flow of If into the chamber was turned on at ∼55 min after the start of exposure of the awake mouse to the sensor and stopped ∼55 min later (see a representative recording from this experiment in Figure 4D). The recorded baseline amplitude declines during anesthesia compared to the pre-anesthesia awake state and takes >60 min to start rising again to return to the pre-anesthesia level during recovery.

### Bidirectional PR in the decapitated head of euthanized mice

In a rodent, neuronal electrical activity ceases <30 s after the last electrocardiographic signal, as indicated by electroencephalographic (EEG) recordings^29^. In dying human patients EEG has been reported to become isoelectric as early as a few minutes before the last heartbeat^30^. However, it is unlikely that any neuron within the brain would still be electrically active hours after complete cessation of cerebral blood flow and perfusion. If neuronal electrical excitation is required for the underlying process that produces a PR then light intensity recording conducted after euthanasia in mice should show the complete disappearance of the PR at a certain latency after the cessation of the heartbeat. The duration of the latency will likely be in the range of 60 – 90 min, given the slowness (see above) of the time course of the effect that originated before death. We performed euthanasia by carbon dioxide inhalation in a total of 8 mice and recorded the PR to 60-min exposures of mice to the light intensity sensor before and after euthanasia. As predicted, a 60-min exposure beginning at 5 min after cessation of the heartbeat showed a decaying PR (Figure 5). The mean peak amplitude of this PR was (280 ± 13, n = 3) au, 54 % of the mean amplitude of the PR before euthanasia induction (531 ± 65 au, n = 3). A 60-min recording from 110 min to 170 showed little or no measurable PR. As in the case of general anesthesia, the half maximal rise time showed a 49% reduction in the postmortem PR (8.1 ± 0.3 min, n = 3) compared to the PR before death (17.2 ± 2.9 min, n = 3). However, when we decapitated the mouse after euthanasia and exposed it and the head and body separately but at the same time to sensor modules of two different instruments, we observed a peri-cranial PR that was initially biphasic (between 5 and 65 min) and then inverted (between 110 and 170, n = 5, Figure 5D). In contrast to the head, the headless body showed a decaying response at the 5-min onset time after clinical death, with the light intensity recording becoming nearly isoelectric at the 110-min postmortem time point. Persistence of the peri-cranial PR at >110 min when there is little or no PR produced by the headless body strongly suggests that the PR primarily, if not exclusively, originates in the brain in mice and passively spreads to the rest of the body where it may be amplified because of its larger volume compared to the head. This prolonged persistence after death also suggests that the source of the peri-cranial PR is unlikely to be neuronal electrical activity.

**Figure 5.**
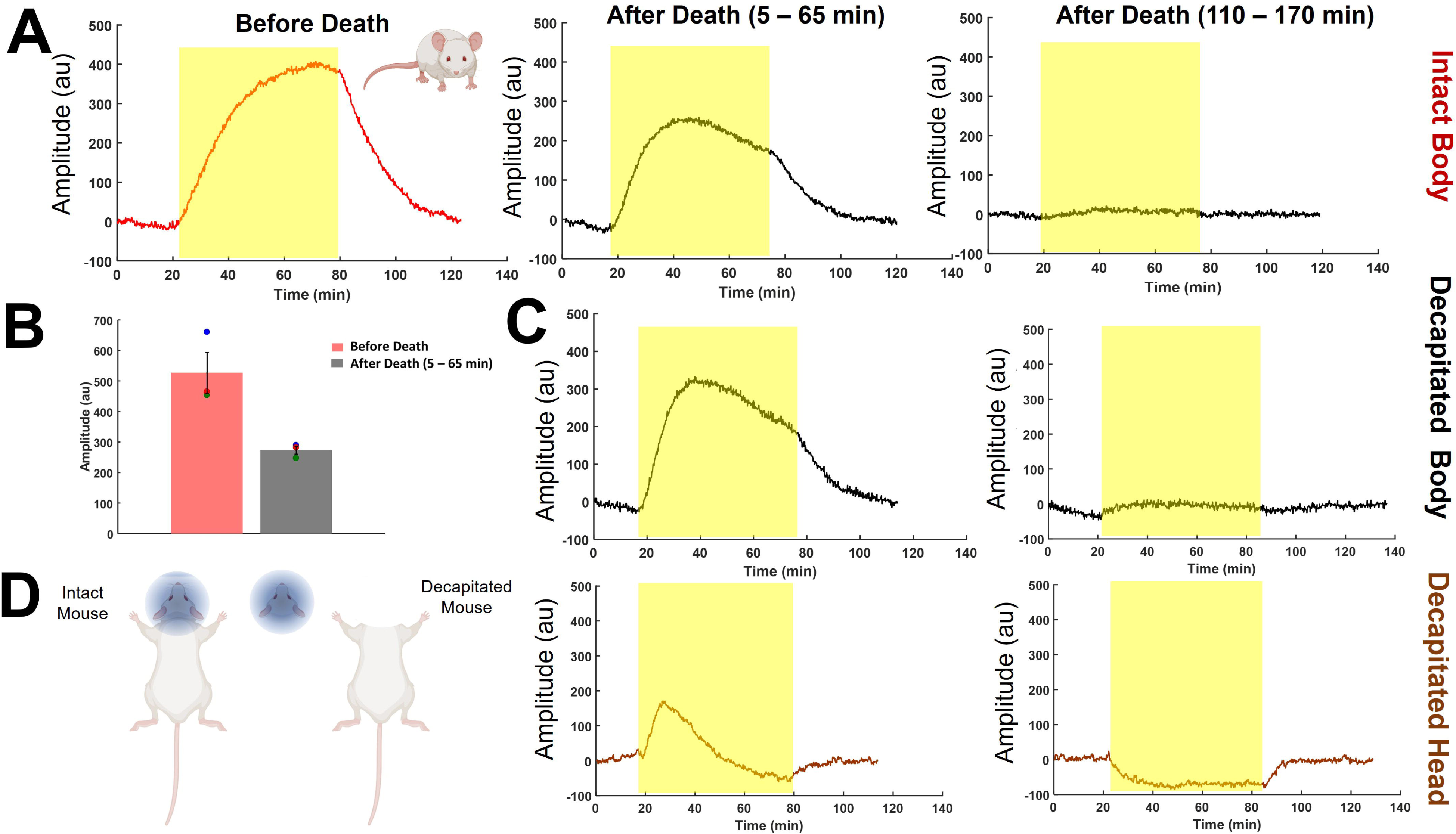
Decline of Peri-somatic PR and Inversion of Peri-cranial PR After Death. **A.** Peri-somatic and peri-cranial PRs in mice before and after euthanasia. The light intensity sensor unit was exposed to the whole body of each mouse at <3 cm in a tube restrainer for 60 min (n = 5). Euthanasia was performed in a carbon dioxide chamber. **B.** The bar plot depicts means and SEMs of 60-min amplitudes before and immediately (5 – 65 min) after death. **C.** and **D.** PRs of the decapitated body (n = 3) and head (n = 3), respectively, at the same two postmortem time points using two different instruments. Images in the figure are obtained from the stock image database of licensed BioRender software.

### Unresponsive patients with primary brain injury show reduced PR and restoration with recovery

To examine whether the amplitude of PR is reduced in unresponsive and clinically presumed unconscious adult patients with primary brain injury due to any cause, we conducted 2 – 3 recordings of 30-min peri-manual PR on different days in each of 3 patients being treated in the ICU. The first patient (a 69-year-old male) had acute encephalopathy supervening on Parkinson’s disease with dementia (PDD). He recovered from the acute condition over several days. The second patient (a 47-year-old male) suffered from severe intracerebral hemorrhage and remained in the same state as determined clinically over the period during which the recordings were conducted. The third patient (a 75-year-old male) had a large subdural hematoma and remained clinically unresponsive and unconscious during our recordings. None of the patients received sedatives. Their results were compared with those obtained from 1 – 3 recordings carried out in 4 conscious healthy adults (2 males and 2 females) under the same conditions in the ICU. The first recordings in all three patients showed significant reduction in mean peak PR amplitude (251 ± 110 au) compared to the mean peak PR amplitude in the healthy controls (667 ± 84 au, n = 4, p = 0.027, Student’s pooled t-test, Figure 6A). Subsequent recordings in each patient are shown in Figure 6B. The first patient showed a progressive increase in peak PR amplitude with clinical recovery, achieving a normal appearing PR 9 days after the initial nearly flat PR. The other two patients showed slight progressive reduction of the peak amplitude, consistent with lack of change in their clinical condition. As seen in the prior recordings in human subjects and mice, in the awake state, PR does not decline until its source (e.g., hand, head or body) is removed from the proximity of the light intensity sensor. By contrast, a noteworthy common feature in all three patients is the premature decline of the PR with or without the occurrence of more than one peak before the removal of the hand. This observation suggests an inability of the injured brain of these patients to maintain a high constant level of consciousness, assuming that the PR amplitude reflects this level. Photo-modulatory data from these patients and healthy controls also allowed us to again test whether PR is simply caused by the change in temperature due to body heat. The PR peak amplitudes ranged from 31.5 au through 758.3 au. However, the corresponding body temperatures measured clinically in patients varied only from 97.6°F through 98.7°F. We correlated the patients’ body temperatures with the highest peak amplitude of their PRs on each day of recording and found that there was no significant correlation between them (r = −0.1, p = 0.74, Pearson’s correlation analysis, Figure 6C), confirming that changes in body temperature cannot account for the changes in PR amplitude.

**Figure 6.**
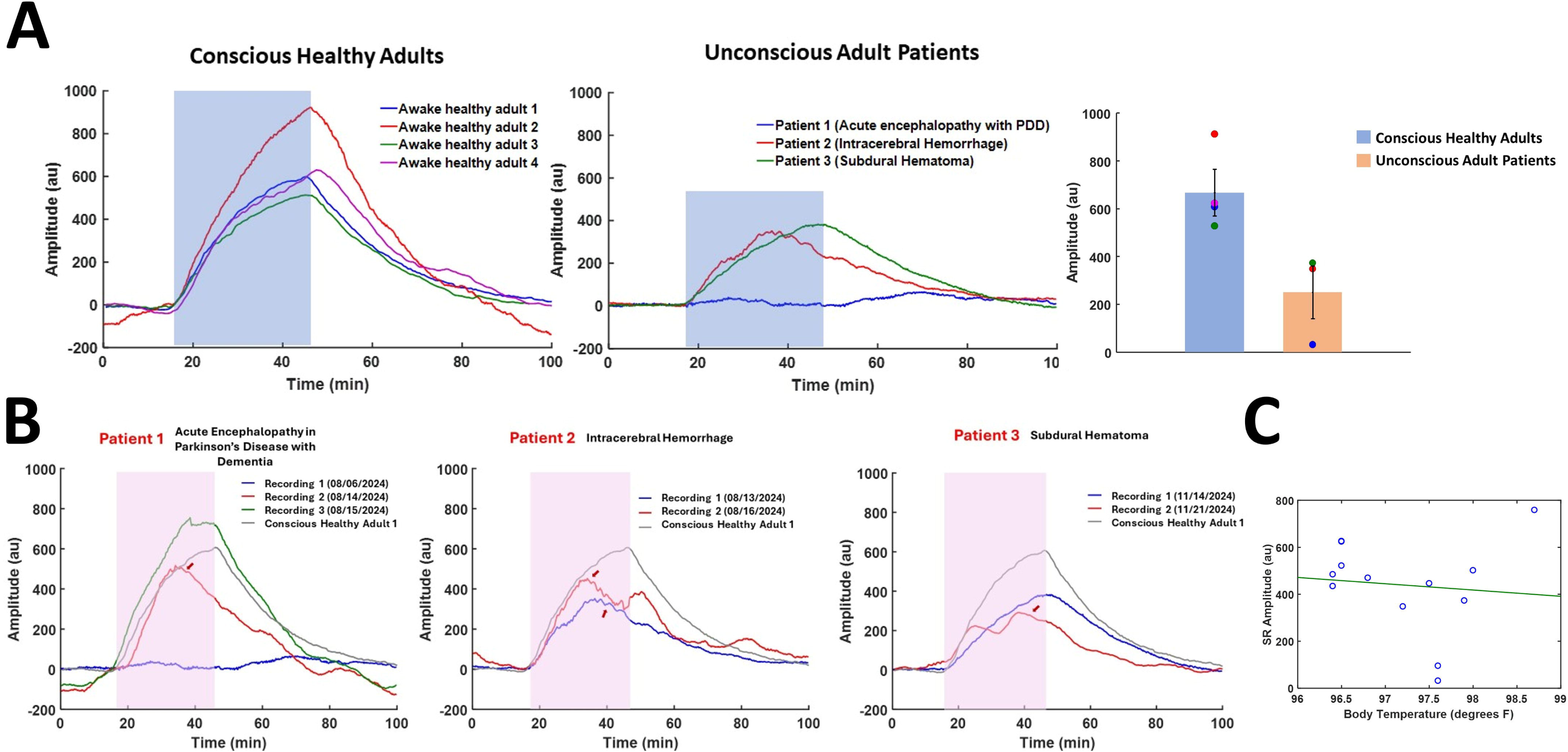
Unconsciousness due to Primary Brain Injury is Associated with Alteration of PR Amplitude and Time Course. **A.** Peri-manual PRs to 30-min exposure to the sensor unit in awake conscious healthy adults (n = 4) and unconscious adult patients with injury to the brain (n = 3) admitted to the intensive care unit (ICU). The bar plot, representing means and SEMs, shows statistically significant ∼72% reduction in the peak amplitude of the PR in patients compared to conscious healthy adults. **B.** Peri-manual PRs obtained on different days during each of the three patients’ stay in the ICU. The short red arrows point to the premature decline of the PR in all three patients before the termination of the peri-manual exposure. The dates of the recordings are indicated in the graph labels. The recording in gray is from the conscious healthy adult 3 from **A**. **C.** A graph of recorded peak PR amplitudes versus body temperatures in all patients and two healthy adults in whom temperature readings were available, showing no correlation between the two variables.

## Discussion

Our experiments demonstrated: first, that light emitted by a low power laser LED when placed near the body of a living organism is correlated with a striking, easily measured, and reproducible and visible change in the digitally recorded intensity of the amount of light that is diffracted. The change takes >20 min to rise to a maximum and, upon moving the instrument sensor away from its source, to decline to the baseline. It is bidirectional, with the direction of change being dependent on the type of organism that is exposed to the instrument sensor, i.e., the intensity decreases in human subjects and mice and increases in several invertebrates. The recorded change is qualitative, and while it can be precisely quantified, it does not require statistical analysis for detection. Second, the amplitude of the PR, representing the amount of change in intensity, declines with distance from the body. Third, two kinds of variations in the level consciousness are associated with qualitative and quantitative changes in PR. General anesthesia in mice is correlated with a faster rise and reduced amplitude of the PR, with recovery resulting in restoration of the rise time, and in some cases also the amplitude, to that seen in the awake state. Furthermore, exposure of a decapitated head from a euthanized mouse to the sensor module of the instrument results in a biphasic PR initially and then an inverted PR that persists for ∼3 hours after death. In ICU patients who are clinically unresponsive and unconscious due to a primary brain injury or impairment, the PR is substantially attenuated and shows a premature decline, despite persistent exposure. In one patient, a virtually absent PR dramatically returned to normal when the patient recovered clinically. These findings provide evidence for a previously unrecognized biophysical effect detectable in the peri-somatic space, which appears to originate in the brain. The effect’s amplitude is modulated by changes in the level of consciousness, with its direction of modulation being determined by the viability of the mammalian brain and its source being invertebrates, as opposed to mammals.

The underlying cause of this newly discovered phenomenon is not clear at present. We have been able to rule out contributions from known factors such as electromagnetic emissions, humidity, breathed air, sound and ultrasound, and heat transfer by conduction and convection from the body. Body heat transfer by radiation cannot also account for PR because of the results of experiments with water bath and the lack of correlation of the large variations in PR amplitude in patients and healthy controls with their body temperature that varied slightly within the normal range. However, given that temperature does have some influence on light intensity recordings, it is possible that a presently unknown classical mechanism analogous to radiative heat transfer is responsible for modulating the intensity of diffracted light waves.

An alternative possibility is that the effect involves a quantum process, which somehow spreads in space.. A third alternative might be a quantum process at the source within neurons spreading spatially through a non-quantum classical process.

## Conclusions

Which of these hypotheses or some other entirely different explanation might account for the photo-modulatory effect await further investigation. But we can at least conclude here that light intensity recordings with the noninvasive photoelectronic instrument developed in our laboratory could serve as a new neurophysiological technique to probe changes that reflect alterations of the level of consciousness in both basic and clinical neuroscience. It is important to note however that the recorded effect may not be specific to consciousness but may reflect a broader cellular or metabolic process.

## Methods

### Description of the Photoelectronic Instrument

We designed and constructed 2 versions of a photoelectronic instrument involving a sensor module containing a low power 5 mW, 650 nm, 3 mm aperture red dot laser light-emitting diode (LED) without an automatic power control circuit, and up to 5 photodiodes. The sensor module in each version is connected to a microcontroller board (STM32, DIANN, China) with a 12-bit analog to digital converter. Version 1 sensor module has a partition containing a vertical double slit (1 mm width slits separated by 2 mm) placed between the laser LED and a horizontal array of 5 photodiodes, which produces an interference pattern. The photodiodes sample the bright interference bands. In version 2, 4 photodiodes form the vertices of a quadrilateral around a central photodiode. The outer photodiodes sample the diffracted light waves emerging from the laser LED.

### Data acquisition and display

The analog channel output from all photodiodes was fed into the analog pins of STM32 and digitized at 12 bits on a 0 – 4095 scale. The digital output sampled at 100 Hz was streamed and logged with a serial monitor program (Serial USB Terminal version 1.55 on Android tablet, Kai Morich; CoolTerm version 2.0.1, Roger Meier on Windows 11 PC). The raw voltage values from the 4 outer channels representing diffracted beams were subjected to principal component analysis to obtain the component of their baseline deviations that is shared across all channels by the highest amount (1st principal component score), which amounted to >97% in all recordings. A further normalization step of this first principal component score time series involved subtraction of its amplitude at each time point from its first time point, which resulted in an inversion of all values. The inversion allowed us to depict the amount of intensity decrease as a positive response whose amplitude represented its strength. The displayed plot of amplitude as a function of time was a smoothed running average of the amplitudes over 1 – 5 s in all figures. All data transformations and plotting were done using scripts written in house using the MATLAB (Mathworks, Inc., Natick, MA, USA) technical computing language.

### Exposure of photo-modulatory sources to the sensor module of the instrument

Procedures involving healthy human volunteers (n = 22) and unresponsive patients (n = 3) were performed under protocols approved by Houston Methodist Research Institute (HMRI) institutional review board (IRB). Informed consent forms signed by the volunteers and the authorized legal representatives of the patients were also approved by the IRB. All procedures in human subjects were performed in accordance with the relevant guidelines and regulations. Experiments involving mice (BALB/c mice obtained from The Jackson Laboratory, Bar Harbor, ME, USA, n = 38) were approved by the HMRI institutional animal care and use committee. In all experiments a 30 – 45 min baseline was recorded before and after each exposure of a photo-modulatory source. All procedures in mice and invertebrates were performed in accordance with the relevant guidelines and regulations. Euthanasia in mice was conducted in accordance with the American Veterinary Medical Association Guidelines for the Euthanasia of Animals (2020). All experiments reported are in accordance with ARRIVE guidelines.

In human subjects, exposure of the head to the sensor module at 1 – 30 cm involved the left or right temple just above the ear. Exposure of the left or right hand involved the palmar surface at similar distances. In the ICU recordings the hand was positioned at 15 cm from the sensor module on a plastic frame during the pre-exposure and post-exposure 45-min baseline recordings. The actual response to exposure of the hand was recorded by placing the hand at 1 cm above the sensor module on the plastic frame. Exposure of mice, or their decapitated heads, was carried out by lightly restraining each live mouse, or placing a dead mouse or head, in a perforated plastic tube with a 4-cm diameter to allow limited front and back, side to side and up and down movement of the mouse and free movement of air. The tube was placed directly over the sensor module, or in the case of the isoflurane (2%) anesthesia experiments, inside the anesthesia chamber placed above it. For injectable anesthetics, the intraperitoneal route was used (sodium pentobarbital 70 mg/kg, propofol 100 mg/kg and ketamine 100 mg/kg xylazine 18 mg/kg). The distance of the mouse from the sensor module was less than 5 cm. 9 blue crabs, 3 crayfish and 6 cherrystone clams were placed one at a time in a closed polypropylene container. The container was then placed over the sensor module. Before exposure of the animal in the container to the sensor module we exposed the empty closed container itself to the sensor module to make sure that it did not induce an PR. Exposure of California blackworms was done by pouring a clump (containing >200 worms) of them along with the water in which they were shipped by the vendor (Carolina Biological Supply Company, Burlington, NC, USA) into a polyethylene bag and holding the bag over the sensor module at 1 cm. The bag with the water alone was exposed to the module as control to make sure it was ineffective in inducing a PR. All recordings were done at room temperature, except in experiments done to measure the effect of temperature.

### Data Analysis

Measured quantitative PR parameters used were mean peak amplitude and half-maximal rise time. Statistical comparison of these parameters between different conditions was done using two-tailed Student’s t tests. In the ICU recordings Pearson’s correlation analysis was done on the PR and body temperature values.

## RESOURCE AVAILABILITY

### Lead contact

Additional information and requests for data and scripts should be directed to the lead contact Dr. Santosh A. Helekar.

### Materials availability

All materials and methods are described in the paper.

### Data and code availability

Data: All data are presented in the paper and supplementary information. Additional information can be requested from the lead contact.

Code: MATLAB scripts mentioned in the paper <u>can</u> be provided upon request by the lead coantact

## ACKNOWLEDGMENTS

The author thanks Blessy John and Alvin Saldon for constructing the photoelectronic instruments, Blessy John, Lisa Nguyen, Shashank Hambarde and Arvind Pandey for assisting with the human and animal experiments, Anh Nguyen for providing access to ICU patients, and Jayant S. Vaidya and Ronald Fisher for extensive discussions on the study experiments and results. This work was supported in part by anonymous philanthropic research funding provided to the author to conduct scientific research on consciousness.

## AUTHOR CONTRIBUTIONS

Study conception and design; data collection, processing and management; analysis and interpretation of results; draft manuscript preparation; and funding acquisition – S. A. H.

