## Supplementary material for "Peri-somatic modulation of diffracted light and its variation with consciousness": Figure S1, Figure S2 and Figure S3

### Supplementary Information

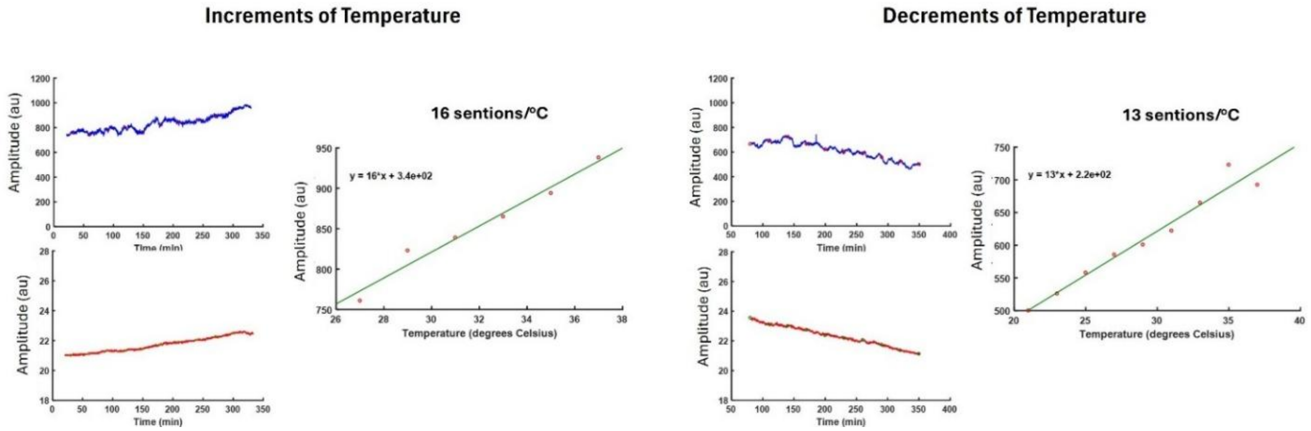


**Fig. S1.**

**Change in Amplitude of the Instrumental Baseline due to Change in Temperature**
**Left –** A beaker containing water was warmed in 2°C increments using an immersion water heater and a temperature regulator while being exposed to the instrument sensor. The recorded changes in the instrumental baseline (top) and water temperature (bottom) and a graph showing significant linear correlation between the two variables are plotted. **Right** – A beaker containing water was cooled in 2°C decrements by dropping ice cubes one at a time in it and similar graphs as on the left are plotted.

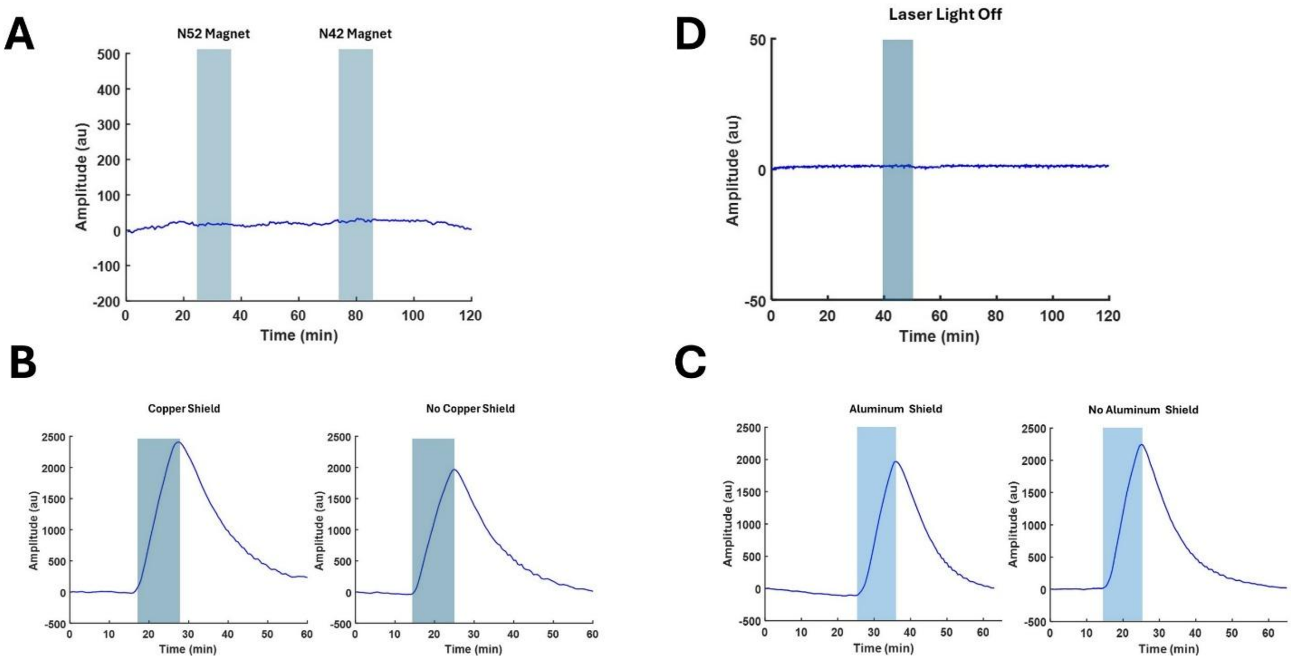


**Fig. S2.**
**Static Electric Fields and Endogenous Ultraweak Photon Emissions do not Contribute to the Photo-modulatory Effect**
**A.** Exposure of N52 and N42 neodymium magnets to the instrument sensor shows no effect on the recorded baseline. **B**. The peri-manual PR amplitude shows no reduction compared to unshielded control when a ~0.6 mm-thick copper shield is placed on the instrument sensor to attempt to obstruct the effect of the hand. **C.** Aluminum (~0.8 mm-thick) barrier also does not block the peri-manual PR. **D.** Light from the laser LED is required for the photodiodes to record the photo-modulatory effect.

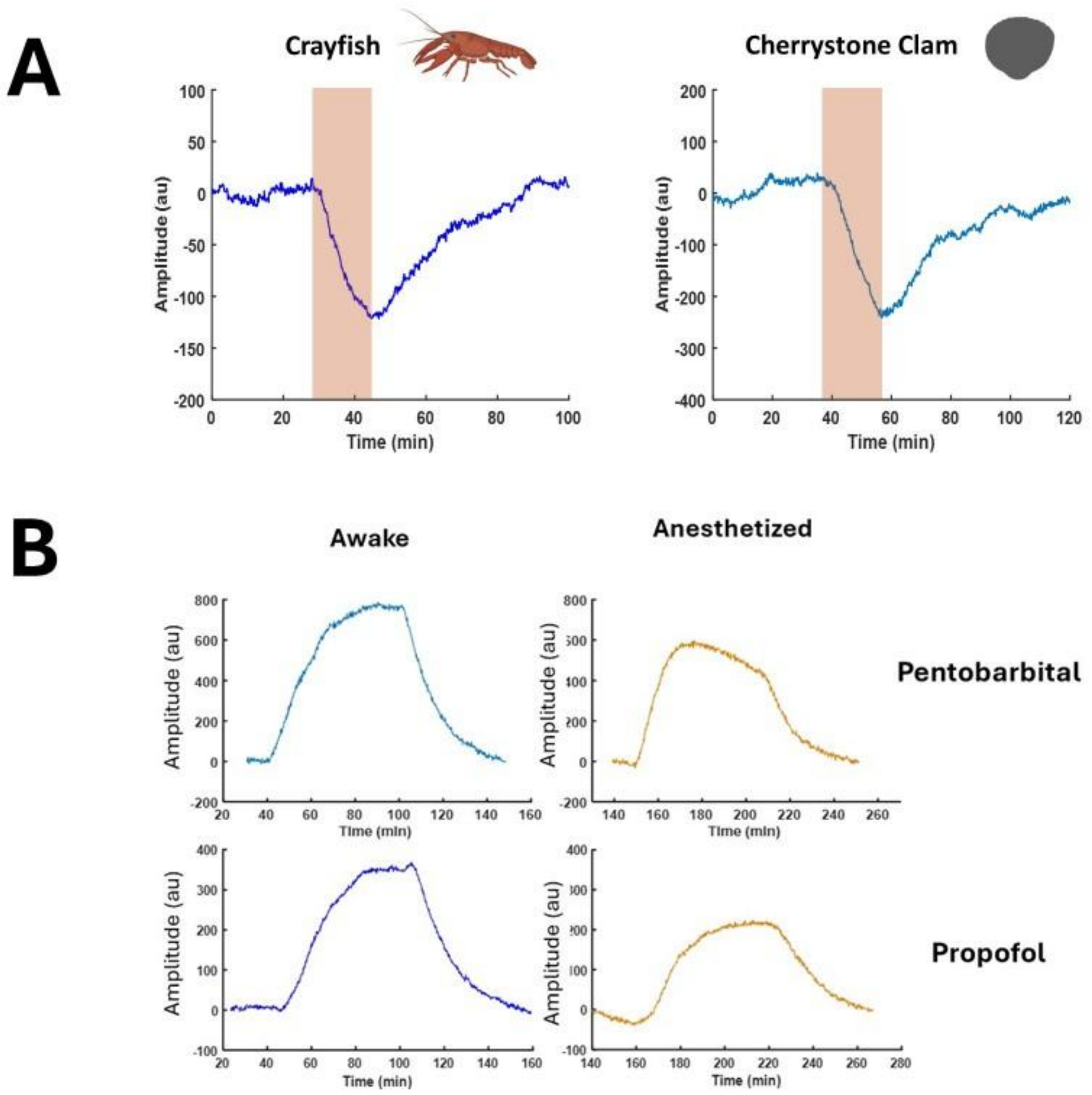


**Fig. S3.**

**PRs in Other Invertebrates and Effect of Other General Anesthetics on the PR in Mice**

**A.** PRs elicited by 15-min exposure of a crayfish and a Cherrystone clam to the instrument sensor **B**. Pentobarbital and propofol produce similar reductions of the PR peak amplitude as those produced by isoflurane and ketamine/xylazine. Images in the figure are obtained from the stock image database of licensed BioRender software.
